# PD-1 expression on NK cells can be related to cytokine stimulation and tissue residency

**DOI:** 10.1101/2021.03.29.437486

**Authors:** Arnika K Wagner, Nadir Kadri, Chris Tibbitt, Koen van de Ven, Sunitha Bagawath-Singh, Denys Oliinyk, Eric LeGresly, Nicole Campbell, Stephanie Trittel, Peggy Riese, Tatyana Sandalova, Adnane Achour, Klas Kärre, Benedict J Chambers

## Abstract

Although PD-1 was shown to be a hallmark of T cells exhaustion, controversial studies have been reported on the role of PD-1 on NK cells. Here, we found by flow cytometry and single cell RNA sequencing analysis that PD-1 can be expressed on MHC class I-deficient tumor-infiltrating NK cells *in vivo*. We also demonstrate distinct alterations in the phenotype of *PD-1*-deficient NK cells which in part could be attributed to a decrease in tumor-infiltrating NK cells in *PD-1*-deficient mice. NK cells from *PD-1*-deficient mice exhibited a more mature phenotype which might reduce their capacity to migrate and kill *in vivo*. Finally, our results demonstrate that PD-L1 molecules in membranes of *PD-1*-deficient NK cells migrate faster than in NK cells from wildtype mice, suggesting that PD-1 and PD-L1 form *cis* interactions with each other on NK cells.

## INTRODUCTION

Natural killer (NK) cells are innate lymphoid cells (ILCs) that can kill tumour cells, stressed or virus-infected cells ^1–3^. NK cell activation is dependent on signals from activating and inhibitory receptors as well as pro-inflammatory cytokines ^4^. Activating NK cell receptors can recognise stress-induced molecules, which induce phosphorylation events that may culminate in the release of cytotoxic granules and cytokines ^5^. Healthy cells are protected from killing by NK cells because of the expression of self-MHC class I molecules (MHC-I) on their surface which act as ligands for dominant inhibitory receptors ^6^. These receptors include killer-cell immunoglobulin-like receptors (KIRs) in humans, Ly49 molecules in mouse and NKG2A in both species ^7^. Engagement of inhibitory receptors results in recruitment of phosphates such as SHP-1, SHP-2 and SHIP-1, and dephosphorylation of signaling molecules which prevents NK cell-mediated killing.

NK cells express also non-MHC-I recognizing inhibitory receptors such as TIGIT, LAG-3, CTLA-4 and PD-1, molecules known as checkpoint receptors. Clinically, antibodies against CTLA-4 and PD-l (or it ligand PD-L1) have been found to be relatively successful in therapy to certain forms of solid cancer ^8,9^. Similar to KIR and Ly49 molecules, checkpoint receptors can recruit and activate phosphatases (DOI: 10.1126/sciadv.aay4458). Several studies have identified subsets of NK cells expressing CTLA-4 ^10^ and PD-1 ^11–14^ in various disease settings but also in healthy individuals ^15^. Furthermore, there is accumulating evidence that NK cells participate in the therapeutic effects of antibodies against PD-1 or PD-L1, expecially against tumors with low MHC-I expression ^11,12,16–21^.

Recently PD-1 expression was detected early in the development of some ILC subsets, which was thought to play a role in the development of ILC responses and ILC subsets could be depleted with anti-PD-1 antibody ^22^. These data raise the question of if and how PD1 is involved in NK cell development and education, and how a chronic lack of PD1 may affect NK cell functions. In the present study, we examined the role of PD-1 in NK cell function using NK cells from *PD-1*-deficient mice as well as the potential role of PD-1/PD-L1 interactions in controlling NK cell activity.

## RESULTS

### NK cell phenotype and population sizes are affected in *PD-1*^−/−^ mice compared to wild-type mice

Although PD-1 plays an important role in the development of ILCs ^22^, no studies to date have examined the phentotype of NK cells from PD-1-deficient mice. Furthermore, lack of PD-1 has been shown to affect T and B cell development, as well as maturation ^23–26^. When comparing the maturation status of NK cells from spleens of wildtype (WT) and *PD-1*^−/−^ mice ^27^, NK cells from *PD-1*^−/−^ mice exhibited an increase in frequency of mature phenotype (CD11b^+^CD27^−^) NK cells (Figure 1a). This appears to take place at the expense of CD11b^+^CD27^+^ NK cells as this subset was reduced in *PD-1*-deficient mice while the size of CD11b^−^CD27^+^ NK cell populations was not affected (Figure 1a). In line with the fact that NK cells derived from *PD-1*-deficient mice exhibited a more mature phenotype, the frequency of KLRG1^+^ NK cell subset^28^ was also increased in *PD-1-*deficient mice compared to WT mice (Figure 1b). In addition, the frequency of CD62L, which is important for NK cell migration ^29^, was reduced in NK cells derived from *PD-1*-deficient mice (Figure 1c).

**Figure 1.**
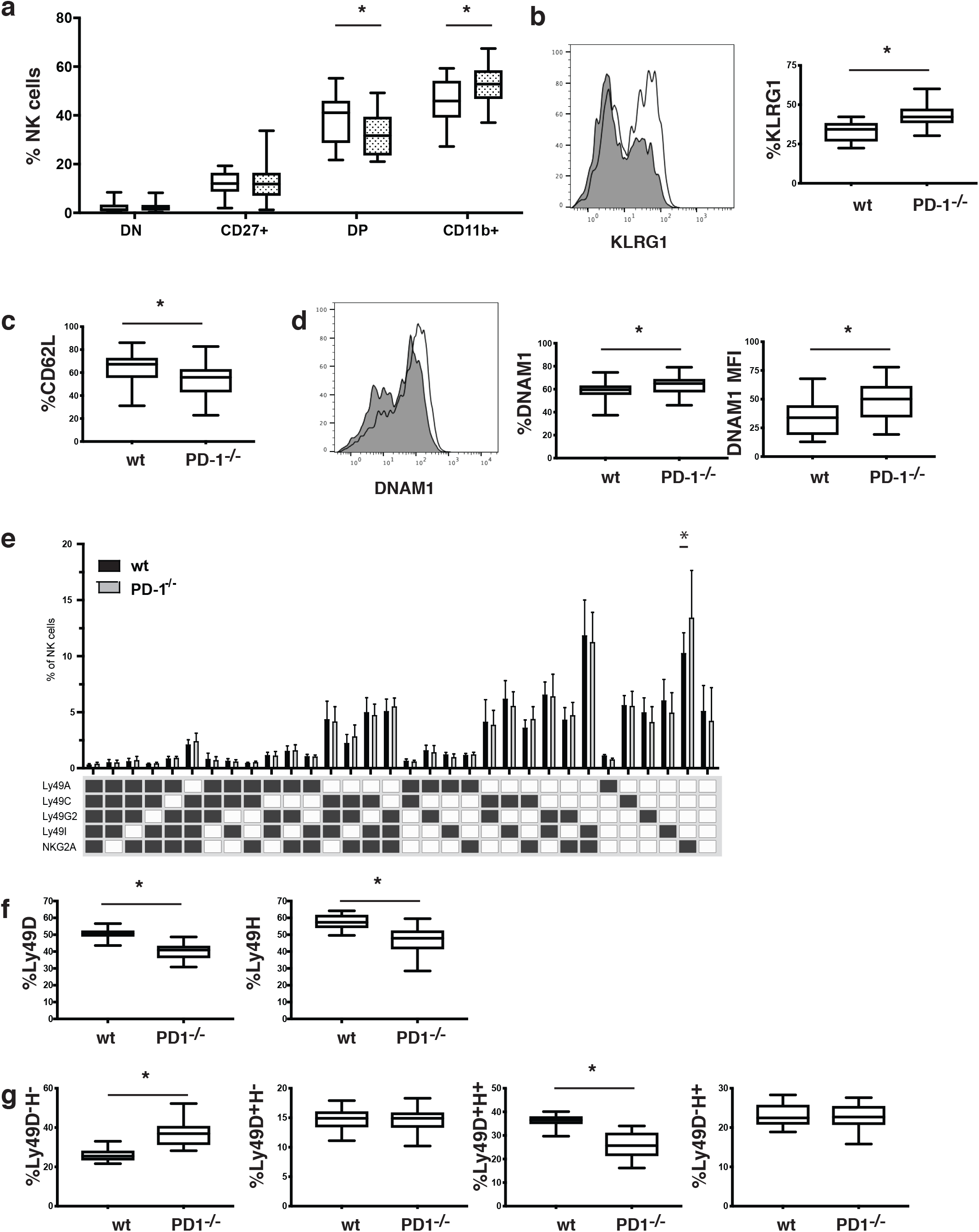
Phenotype of NK cells from WT and PD-1^−/−^ mice. (a) Expression of CD11b and CD27 on NK cells from WT (open boxplots) and *PD-1*^−/−^ (shaded boxplots) mice (*p<0.01 Mann-Whitney test, n=18-20 mice (b) Expression of KLRG1 on NK cells from WT and PD-1^−/−^ mice (*p<0.01 Mann-Whitney test, n=18-20 mice). (c) Expression of CD62L on NK cells from WT and PD-1^−/−^ mice. (d) Expression of DNAM-1 on NK cells from WT and PD-1^−/−^ mice, bar graphs represent percent expressing cells and the mean fluorescent intensity of expression (*p<0.01 Mann-Whitney test, n=18-20 mice. (e) Expression of inhibitory Ly49 molecules and NKG2A on NK cells from WT and PD-1^−/−^ mice (*p<0.01 Mann Whitney n=18-20 mice). (f) Expression of activating Ly49 molecules on NK cells from WT and PD-1^−/−^ mice. (*p<0.01 Mann-Whitney test, n=18-20 mice). (g) Expression of Ly49D and Ly49H populations on NK cells from WT and PD-1^−/−^ mice (*p<0.01 Mann-Whitney test, n=18-20 mice).

It has recently been shown that PD-1 affects DNAM-1 expression on CD8 T cells ^30^. We observed an increased frequency of DNAM-1^high^ NK cells as well as increased expression levels of DNAM-1 in *PD-1-*deficient mice compared to WT mice (Figure 1d). This confirms that PD-1 can modulate DNAM-1 expression not only on CD8 T cells but also on NK cells.

We further analyzed expression of the inhibitory receptors on the NK cells^31^, and compared the repertoire of inhibitory molecules on NK cells from WT and *PD-1*-deficient mice. We did not find any major differences in Ly49 and NKG2A expression between these mice, apart from an increase in the NKG2A^single^ population on NK cells from *PD-1*^−/−^ mice (Figure 1e). The frequency of the activating Ly49D and Ly49H molecules was reduced in *PD-1*^−/−^ mice, and this appeared to be due to a reduction in the frequency of the Ly49D^+^Ly49H^+^ NK cell population (Figure 1f and g). The expression levels of other activating receptors for example NKG2D and CD244 were not significantly different between WT and *PD-1*-deficient mice (Supplemental Figure 1a-b).

Lack of PD-1 has been associated with the accumulation of exhausted T cells^24^. Lag3, CD39 and TIGIT are can be used as markers for T cell exhaustion^24^. Comparing the NK cells from WT and *PD-1*^−/−^ mice, we observed only a small changes in the frequency of CD39^+^ NK cells in *PD-1*^−/−^ mice and the frequency of LAG3^+^ NK cells (Supplemental Figure 1c-d). In addition, the surface expression of PD-L1, the ligand for PD-1, was not significantly different between the two mouse strains (Supplemental Figure 1e). In addition, we did not observe any difference in the expression of GITR, CXCR3 or CXCR4 (Supplemental Figure 1f-h).

We were concerned that some of the phenotypic changes that we observed on *PD-1-*deficient NK cells might be due to perturbations caused by T cells lacking PD-1 ^32^. Therefore we compared NK cells from *PD-1xRAG1*^−/−^ and *RAG1*^−/−^ mice since both these mice have neither T nor B cells. Similarly to T and and B cell-competent mice, NK cell maturation was still skewed in *PD-1xRAG1*^−/−^ mice with increased frequencies of CD11b^+^CD27^−^ and KLRG1^+^ NK cells compared to *RAG1*^−/−^ mice (Supplemental Figure 2a-b). However, we no longer observed any significant difference in the frequency of CD62L^+^ NK cells between *RAG1*^−/−^ and *PD-1xRAG1*^−/−^ mice (Supplemental Figure 2c).

DNAM-1 expression levels were increased still on NK cells from *PD-1xRAG1*^−/−^ mice but unlike in T and B cell competent mice, the frequency of CD39 expressing NK cells was increased in the *PD-1xRAG1*^−/−^ mice (Supplemental Figure 2d and 2e). In contrast to the *PD-1^−/−^* mice, analysis of the expression levels of inhibitory receptors revealed no longer any difference in frequency of the NKG2A^single^ NK cell population between *RAG1*^−/−^ and *PD-1xRAG1*^−/−^ mice (Supplemental Figure 2f).

While the frequency of Ly49D^+^ NK cells was reduced in *PD-1xRAG1*^−/−^ mice, there was no difference in Ly49H expression between *RAG1*^−/−^ and *PD-1xRAG1*^−/−^ mice. The reduction in the Ly49D population appeared to be occuring mostly in the Ly49D^+^Ly49H^−^ subset and not in the Ly49D^+^Ly49H^+^ population (Supplemental Figure 2g and 2h).

In summary, we observed in mice lacking PD-1 increased NK cell maturation combined with higher DNAM-1, KLRG1 expression and reduced Ly49D expression.

### Elimination of MHC-I-deficient cells is impaired in *PD-1*^−/−^ mice

Chronic loss of PD-1 could potentially affect not only the phenotype of NK cells as outlined above, but also their function. The recognition and elimination of cells expressing reduced MHC-I levels is a hallmark of NK cell function and education ^33–35^. We therefore examined the ability of *PD-1*-deficient and WT mice to eradicate MHC-I^neg^ spleen cells. We observed a significant reduction in the ability of *PD-1*^−/−^ mice to eliminate MHC-I^neg^ splenocytes compared to WT mice (Figure 2a). However this impairment was not at the level seen in MHC-I^−/−^ mice, suggesting that NK cells might be affected by non-MHC-I factors such as increased maturity of NK cell populations as described above.

**Figure 2.**
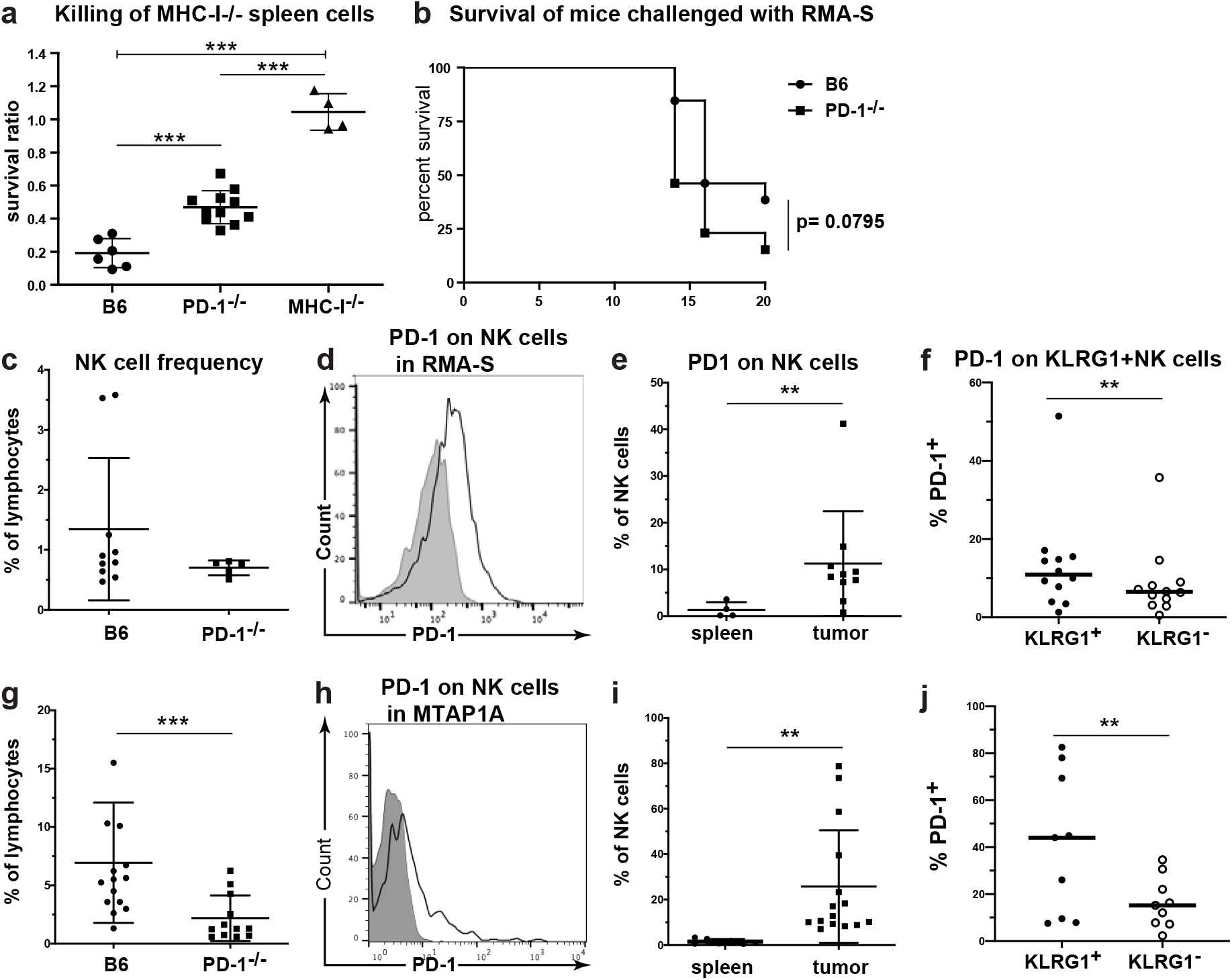
*PD-1-*deficient mice exhibit poor rejection of MHC-I-deficient cells. (a) WT and *MHC-I*^−/−^ splenocytes labelled with CFSE were injected and the rejection ratio measured in WT, *PD-1*^−/−^ and *MHC-I*^−/−^ mice ***p<0.001 (ANOVA three separate experiments total of 6-8 mice). (b) Rejection of RMA-S cells injected s.c. WT and *PD-1*^−/−^ mice were given an LD_50_ dose of RMA-S cells (10^5^ cells) and the survival rate of mice was measured (survival measured using log rank test, three separate experiments with n= 15-16 mice). (c) Percent intratumoral NK cells amongst the lymphocyte population in mice receiving RMA-S. (d-e) Expression of PD-1 on intratumoral NK cells in RMA-S treated mice compared to expression on splenocytes (**p<0.001 Mann-Whitney test). (f) Expression of PD-1 on intratumoral KLRG1^+^ and KLRG1^−^ NK cell populations (p<0.05 paired t-test). (g) Frequency of intratumoral NK cells amongst lymphocytes in mice receiving MTAP1A (**p<0.01 Mann-Whitney test). (h-i) Expression of PD-1 on intratumoral NK cells in MTAP1A treated mice compared to expression on splenocytes (**p<0.01 Mann-Whitney test, n=12-15 mice). (j) Expression of PD-1 on intratumoral KLRG1^+^ and KLRG1^−^ NK cell populations in MTAP1A tumors (**p<0.01 paired t-test n=8).

It has been previously demonstrated that anti-PD-1 treatment increases NK cell elimination of MHC-I^neg^ PD-L1^+^ tumors ^12^. To determine if *PD-1*^−/−^ mice also had reduced erradication of a MHC-I^neg^ tumor with low expression of PD-L1^12^, mice were injected with an LD_50_ dose of TAP-deficient RMA-S lymphoma cells. While the survival rate of WT mice was 45%, only 30% of *PD-1*-deficient mice survived (Figure 2b). Comparison of tumor infiltrating NK cells from WT and *PD-1*^−/−^ mice revealed a reduced frequency of tumor infiltration in *PD-1*-deficient mice (Figure 2c). PD-1 was heterogeneously expressed on NK cells infiltrating RMA-S in WT mice (Figure 2d), while splenic NK cells from the same mice exhibited little or no PD-1 expression (Figure 2e). While these findings are similar to previous studies^12^, the frequency of PD-1 expression on the NK cells from our study were significantly lower ^12^.

Next, we examined tumor infitrating NK cells from a second MHC-I^low^ tumor cell line MTAP1A, which is a fibrosarcoma generated from the skin of a *Tap1*-deficient mouse^36^. MTAP1A has low expresssion of PD-L1, and does not express PD-L2 (Supplemental Figure 3). Here again, we found reduced infiltration of NK cells in *PD-1*^−/−^ mice but increased expression of PD-1 on tumor-infiltrating NK cells in WT mice compared to splenic NK cells (Fig. 2g-i and Supplemental Figure 4).

In addition, PD-1^+^ tumor-infiltrating NK cells also displayed increased expression of KLRG1 compared to PD-1^neg^ NK cells (Figure 2f and j). This was observed for tumor-infiltrating NK cells in both RMA-S and MTAP1A, and suggested that PD-1-expressing NK cells might have a more mature phenotype.

### Single cell RNA-seq reveals tissue-specific transcriptional imprinting of tumor infiltrating NK cells

Since it has been suggested that NK cells may express PD-1 through trogocytosis ^37^ and since we observed differences in the phenotype of NK cells from WT and *PD-1*^−/−^ mice, we performed single cell RNA-sequencing (scRNA-SEQ) using the SMART-SEQ2 platform ^38^ on tumor-infiltrating NK cells from mice inoculated with the MTAP1A tumor. We chose MTAP1A over RMA-S since this tumor model gave consistently higher frequency of PD-1-expressing NK cells. SMART-SEQ2 libraries of sorted NK cells were generated from pooled tumors from either WT or PD-1-deficient mice (Supplemental Figure 5a-b). These libraries were filtered and a combined analysis was performed using Seurat v3 ^39,40^ for a total of 371 WT and 375 *PD-1*^−/−^ NK cells after quality control (Supplemental Figure 5c). Outliers expressing very few or very many genes were omitted, as were cells with a high frequency of apoptotic genes. Cells were clustered and projected using UMAP, which delineated five clusters with both WT and *PD-1*^−/−^ NK cells found in all clusters although *PD-1-*deficient cells were over-represented in Clusters 3 and 4. (Figure 3a-c). Differentially expressed (DE) genes were deciphered between all clusters and the top 10 genes per cluster shown by heatmap (Supplemental Figure 6a). Selected genes were plotted using the Violin plot function revealing significantly over-expressed genes in each cluster. Within clusters 3 and 4, we could detect *Pdcd1* (PD-1) transcripts in both WT and *PD-1*^−/−^ NK cell populations, suggesting an active upregulation of *Pdcd1* at the transcriptional level. (Figure 3d). Detection of *Pdcd1* transcript in *PD-1*^−/−^ mice reflects that these mice do not have a complete gene defect but rather a deletion spanning exon 3 and exon 4 of the P*dcd1* gene that prevents protein expression^26^. Our analysis highlighted the heterogeneity of *in vivo* NK responses with distinct patterns of *Prf1* and *Gzma*, *Gzmb* and *Gzmc* expression (Figure 3e). Interestingly, expression of the tissue residency marker *Cd69* was closely aligned with *Gzmc* detection (Figure 3e and g).

**Figure 3.**
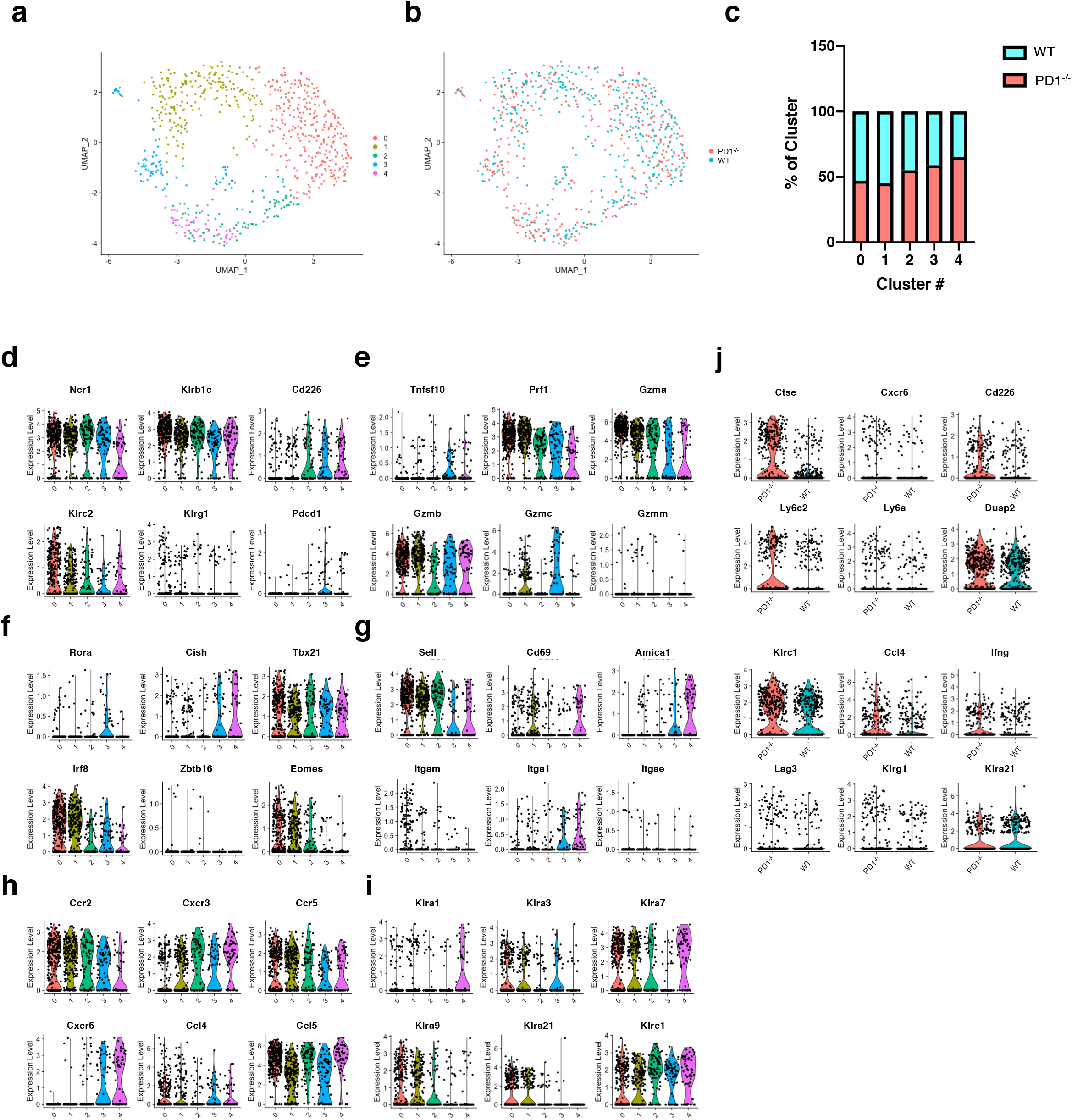
Single cell analysis of intratumoral NK cells. (a, b) UMAP projection of 746 tumor-infiltrating NK cells (371 WT cells, 375 PD-1^−/−^ cells). (c) Percentage of each cluster derived from either WT or PD-1^−/−^ NK cells. (d-i) Violin plots for several genes enriched across various clusters. (j) Violin plots depicting expression of several genes differentially expressed between WT and PD-1^−/−^ NK cells.

Clusters 3 and 4 were defined by a paucity of *Eomes* and *Irf8* whilst being enriched for expression of *Tnfsf10* (TRAIL), *Cxcr6* and *Itga1* (CD49a) (Figure 3e-g). These clusters also had greater expression of *Amica1* (JAML), *Ly6a*, *Il7r* and *Il21r* and lower levels of *Sell* (CD62L) (Figure 3g and Supplemental Figure 6b). Cells found in cluster 3 also had significantly enhanced *Lag3* levels suggesting this population may potentially harbor exhausted NK cells (Supplemental Figure 6b). Taken together these findings indicate that clusters 3 and 4 might represent a more mature/exhausted population and/or a tissue-resident-like subset of NK cells. Finally, we observed differential expression of transcripts for the inhibitory Ly49 genes *Klra1*, *Klra3*, *Klra7* and *Klra9* amongst the different clusters. In particular, Klra3 (Ly49C) seemed to be present in cluster 3 but *Klra1* (Ly49A), *Klra7* (Ly49G2) and *Klra9 (Ly49I)* seemed to be under-represented in the same cluster (Figure 3i). This indicated that different Ly49 subsets of NK cells in conjunction with PD-1 might play a role in the tumor environment as previously seen in other studies ^12^ (Figure 3i).

Comparison of all WT with all *PD-1*^−/−^ NK cells independent of cluster identity determined a total of 54 genes that were significantly altered between these two NK cell populations (Figure 3j and supplemental table I). Amongst those over represented in *PD-1*^−/−^ NK cells were transcripts for *Cd226* (encoding for DNAM-1), *Klrc1* (NKG2A) and *Klrg1*, which is very well in line with our flow cytometry data on spleen NK cells. Furthermore, we found that *PD-1*-deficient NK cells had altered levels of expression for *Cxcr6* and select members of both *Ly6* families, suggesting that NK cells from the *PD-1*^−/−^ mice had a more tissue resident phenotype. *Ifng* and *Ccl4* transcripts were also more abundant in NK cells from *PD-1*^−/−^ mice indicating an influence of PD-1 on *in vivo* NK cell responses (Figure 3j).

### PD-1 is induced on the surface of NK cells after stimulation with cytokines

Since the expression profile of NK cells expressing PD-1 and CXCR6 suggested that they may be connected, we stimulated DX5^+^-enriched NK cells from WT mice with a combinatin of IL-12/15/18 cytokines for 96 hours, which has previously been show to induce memory NK cells ^41^ as well as CXCR6 on the surface of NK cells ^42^. This cytokine stimulation resulted in approximately 10% of WT NK cells expressing PD-1 (Figure 4a). Similar patterns of staining were seen in cytokine-stimulated NK cells from *RAG1*^−/−^ mice (Figure 4b), which ruled out that expression of PD-1 might be on a T cell subset with low CD3 expression, that T cells could induce PD-1 on NK cells or that PD-1 expression was through trogocytosis from T cells.

**Figure 4.**
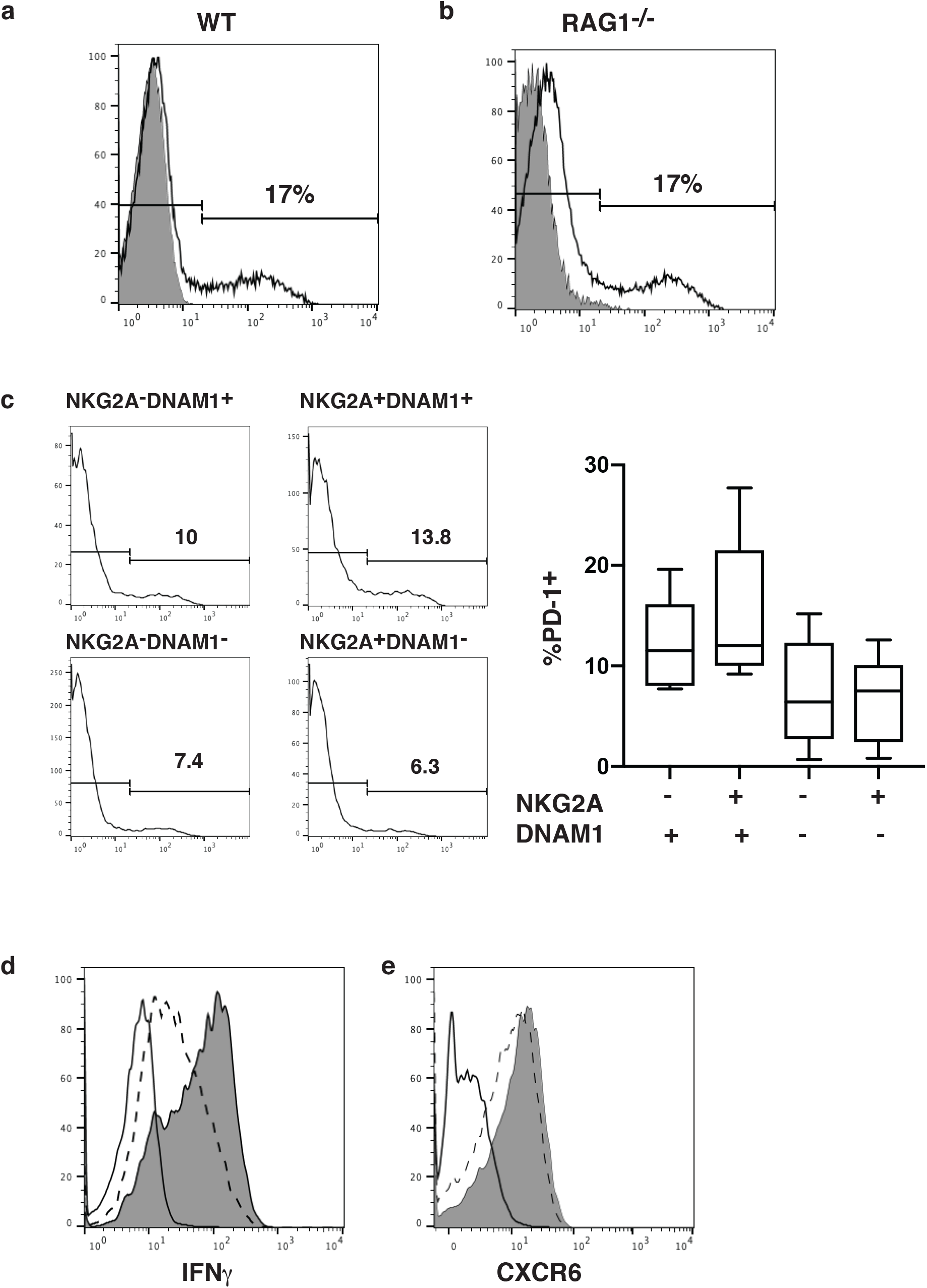
Induction of PD-1 on NK cells following cytokines stimulation. (a) PD-1 expression on WT NK cells (white histogram) following stimulation with IL-12/15/18 for 4 days, expression on PD-1^−/−^ NK cells is shown as comparison (grey histogram). (b) Expression of PD-1 on NK cells from RAG1^−/−^ mouse following cytokine stimulation. (c) Expression of PD-1 on DNAM-1 and NKG2A populations following cytokine stimulation (from four experiments. (d) Intracellular levels of IFNγ following cytokine stimulation of NK cells from (dashed line) WT mice and (shaded line) PD-1^−/−^ mice (representative plot from three experiments). (e) Expression of CXCR6 on the surface following cytokine stimulation of of NK cells from (dashed line) WT mice and (shaded line) PD-1^−/−^ mice.

We have previously demonstrated that we could define functional NK cell subsets based on DNAM-1 and NKG2A expression ^43^. Therefore, we here investigated whether PD-1 expression was confined to a specific NK cell subtype following cytokine stimulation. We assessed the expression of PD-1 on DNAM-1^+^NKG2A^+^, DNAM-1^+^NKG2A^−^, DNAM-1^−^NKG2A^+^ and DNAM-1^−^NKG2A^−^ NK cell subsets. Although we did not observe any significant difference in the percentage of PD-1^+^ NK cells between these subsets, DNAM-1^+^ NK cells had on average an increased percentage of PD-1 expressing NK cells following cytokine stimulation (Figure 4c).

Since the intratumoral NK cells from *PD-1*^−/−^ mice exhibited a trend towards expressing more IFNγ, we examined the intracellular levels of IFNγ in the IL-12/15/18 cytokine stimulated NK cells. We found increased levels of intracellular IFNγ in the *PD-1*^−/−^ NK cells compared to NK cells from WT mice (Figure 4d), confirming that lack of PD-1 might predispose NK cells to increased IFNγ expression. Furthermore, we also found that CXCR6 levels were increased on the surface of NK cells from *PD-1*^−/−^ mice following cytokine stimulation, suggesting a role for PD-1 in controling the expression CXCR6 (Fig 4e).

### PD-1 can form *cis* interactions with PD-L1 on NK cells

The tumor cells used in our experiments had little or no PD-L1 expression. Therefore, our observations that the PD-1 expressing cells is increased on intratumoral NK cells and IL-12/15/18 stimulated NK cells might mean that PD-L1 could interact with PD-1 on NK cells both in *cis* or *trans*. It has been shown previously that inhibitory MHC-class I binding molecules on NK cells could form *cis*-interactions with their ligands ^44,45^. Furthermore, PD-1 and PD-L1 have recently been shown to form *cis*-interactions in artifical lipid structures and in antigen-presenting cells (APCs) ^46^. We therefore assessed whether the movement of PD-L1 was restricted in the presence of PD-1 and determined PD-L1 diffusion on the membranes of NK cells lacking PD-1 compared to WT NK cells using fluorescence correlation spectroscopy (FCS), a method that detects diffusion of molecules and has previously been used to measure the diffusion of receptors in the membrane of NK cells ^45,47^. FCS measurements were performed on the cell membrane, and a series of autocorrelation curves were generated and fitted to the 2D diffusion FCS curve fitting equation. Representative autocorrelation curves with 2D curve fit are shown in Figure 5a. Interestingly, PD-L1 diffused significantly faster on the membrane of NK cells lacking PD-1, compared to PD-1^+^ NK cells from WT mice (Figure 5a and b). Furthermore, we observed a trend for high levels of PD-L1 molecules per μm^2^ on the surface of NK cells lacking PD-1 (Figure 5c). Since molecule crowding factor is ruled out on PD-1^+^ NK cells, the slow diffusion of PD-L1 molecules on cell membrane can be due to specific interactions or clustering. To investigate whether PD-1 and PD-L1 form clusters on the surface of NK cells, the brightness of PD-L1 was quantified that is measured in terms of counts per molecule, diffusing within the observation volume. We observed a tendency towards larger clusters, as the brightness of PD-L1 on PD-1 positive NK cells was higher compared to *PD-1*^−/−^ NK cells (Figure 5d). These data suggest that the PD-L1 on PD-1^+^ NK cells clusters with PD-1, indicating *cis* interactions on the membrane of NK cells. In conclusion, the PD-L1 diffusion faster without any hinders on *PD-1*^−/−^ NK cells whereas in presence of PD-1 on cell membrane PD-L1 diffuse slower, this suggest that PD-L1 might be clustering in *cis* with PD-1 on the cell membrane (Figure 5e).

**Figure 5.**
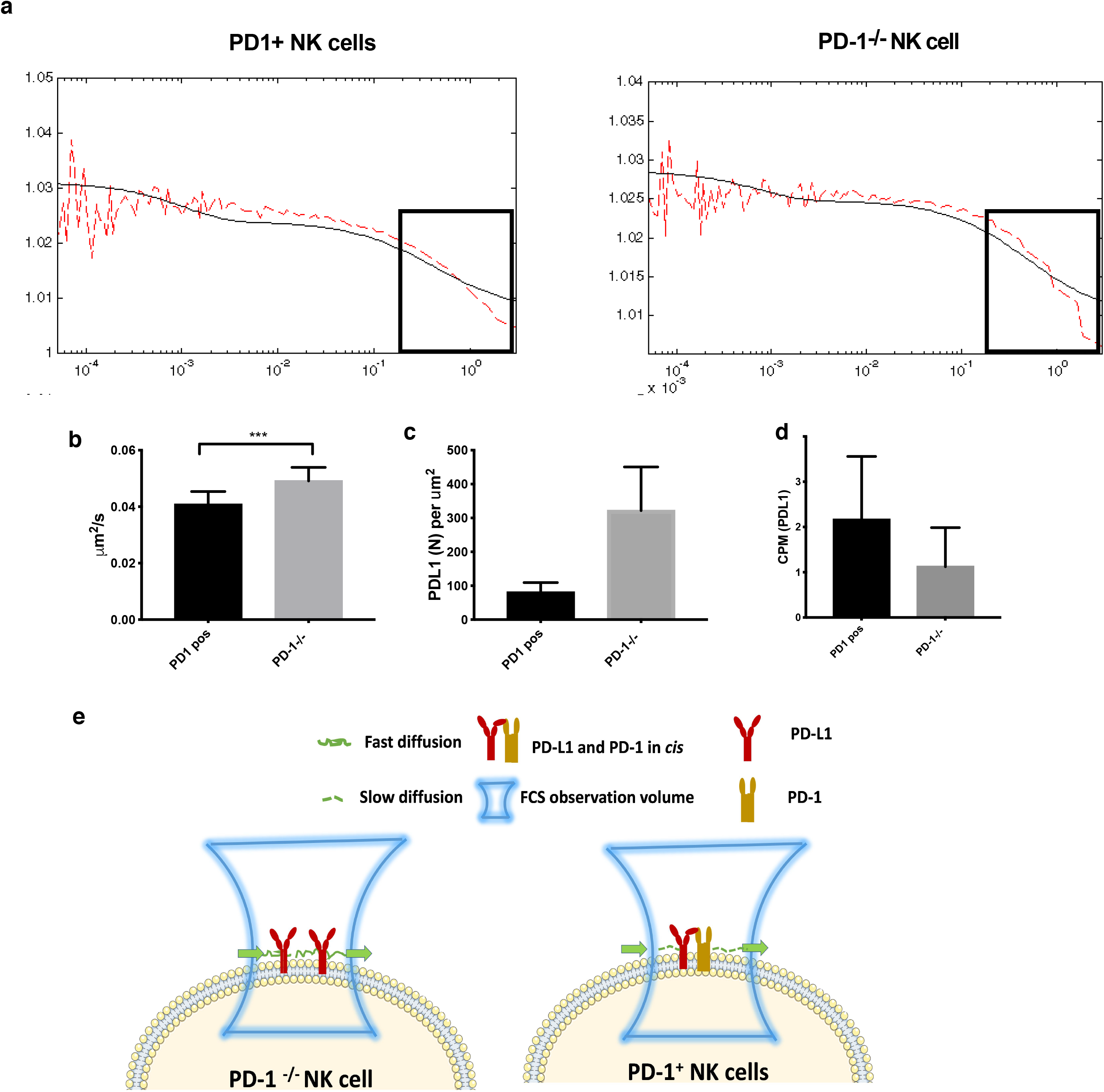
Movement of PD-L1 in WT and PD-1^−/−^ NK cells. (a) Representative FCS auto correlation curves of PD-L1 on PD-1 positive (*right panel*) and PD-1^−/−^ NK cells (*left panel*), decline part of the curve indicates the rate of diffusion on cell membrane. FCS readouts of PD-L1 molecule on PD-1 positive and negative NK cells. (b) The diffusion rate of PD-L1, (c) the density of PD-L1 and (d) the counts per molecule (CPM) of PD-L1 were measured on individual PD-1^+^ NK cells from *RAG1*^−/−^ mice and NK cells from *PD-1xRAG1*^−/−^ mice. (b) The diffusion rate of PD-L1 is faster in the absence of PD-1. (c) The density of PD-L1 was higher on *PD-1xRAG1*^−/−^ NK cells, while (d) the CPM, indicates the size of the cluster measured based on the brightness or number of molecules per entity, PD-L1 clusters was higher when PD-1 was present. (e) Model for PD-L1 movement in the membrane in the presence and absence of PD-1. Comparison between two unpaired groups of 12 NK cells by Mann Whitney test, *** p< 0.001.

Three-dimensional molecular models of the full-length extracellular domains of PD-1 and PD-L1 reveal that their structural features easily allow for the formation of cis-interactions. Indeed, a model of the stalk region of PD-1 (comprising the stretch of residues R147-V170) in extended conformation demonstrates that its length is sufficient to allow both *cis*- and *trans-*interactions with the N-terminal domain of PD-L1 (Figure 6). Our molecular models thus suggest a binding in which PD-1 “tip-toes” to reach PD-L1 with an extended stalk, while keeping the same PD-1/PD-L1 “cheek-to-cheek” interface found in previous crystal structures (Figure 6).

**Figure 6.**
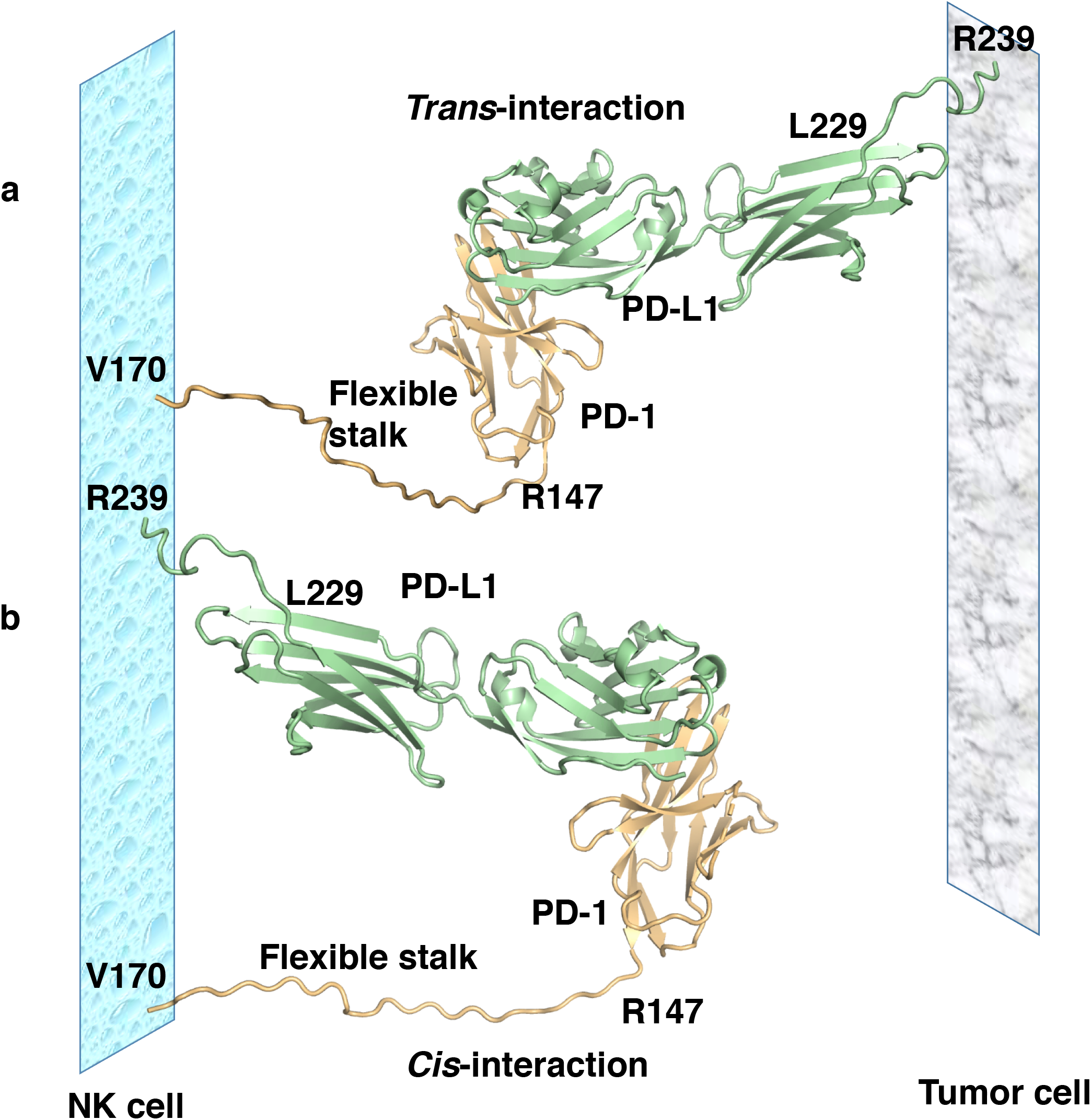
Molecular Modelling of PD-1-PD-L1 *cis*-interaction. The 24 amino residues long stalk region of PD-1 is long and flexible enough to allow for both *trans*- and *cis*-interaction between PD-1 and PD-L1. (a) *Trans*-interaction between PD-L1 on tumor cells and PD-1 on NK cells. (b) *Cis*-interaction between PD-1 and PD-L1 on NK cells. The interactions and mode of binding between the N-terminal part of PD-L1 and PD-1 could be highly similar, as found in the crystal structure of the human PD-1/PD-L1 complex ^65^. The stalk-region of PD-1 (residues R147-V170) was modelled in an arbitrary extended conformation to show that its length is sufficient to allow for *cis*-interaction with the N-terminal domain of PD-L1.

## DISCUSSION

Expression of PD-1 on NK cells has been observed in many human and mouse studies^11,12,14,16,32,48–50^. However, some recent studies suggested that NK cells do not express PD-1 and expression may due to artifact of flow cytometry staining or through interactions with PD-L1 and the NK cells acquiring PD-1 via trogocytosis^37^. However in the present study we could find transcript and surface expression of PD-1 in tumor infiltrating NK cells, as well as upon IL-12/15/18 stimulation of NK cells in culture. Furthermore, we also found that PD-1 was induced on tumor-infiltrating NK cells even though the tumors used in our study expressed little or no PD-L1. NK cells from mice lacking PD-1 displayed phenotypic differences compared to NK cells from WT mice, suggesting that background low levels of PD-1 might still play a role in NK cell homeostasis or in NK cell development. In particular, NK cells from *PD-1-*deficient mice exhibited increased maturation as well as increase in expression of CD226.

We found that *PD-1*-deficient mice were poor at rejecting MHC-I^−/−^ cells and had low NK cell infiltration into tumors expressing low levels of MHC-I *in vivo*. In part, this might be due to the increased maturity of NK cells in *PD-1*-deficient mice, but the reduced frequency of tumor infiltrating NK cells could also be due to (i) reduced CD62L found on the PD-1^−/−^ NK cells or (ii) reduced survival once these NK cells encounter tumor cells. It is unclear if the increased NK cell maturation is a direct effect on NK cells since we observe very little or no PD-1 on NK cells in circulation. However, others have shown that lack of PD-1 on dendritic cells leads to increased IL-12 and TNF production by dendritic cells ^51^. Thus lack of PD-1 on DCs could indirectly affect NK cell maturation. Furthermore, absence of PD-1 on T and B cells could affect NK cells indirectly as well ^52,53^. For this reason, we crossed *PD-1-*deficient mice to mice deficient in *RAG1*. In the *PD-1xRAG1*^−/−^ mice, we still had more mature NK cells and increased expression of DNAM-1 suggesting that the *PD-1*^−/−^ T and B cells had little effect on the NK cell phenotype. *PD-1*^−/−^ NK cells stimulated with IL-12/15/18 had increased numbers of IFNγ-producing cells suggesting that increased IL-12 from accessory cells in *PD-1*^−/−^ mice^51^ might already prime NK cells to make more IFNγ. Chronic infection and IL-18 expression have previously been associated with higher expression of PD-1 on NK cells ^32,50,54,55^. Even though PD-1 expression on T cells has been associated with exhaustion, it is more likely that it is also a marker for activation and that its expression controls T cells from being overly activated ^24,56^. Thus, PD-1 expression on NK cells might play a similar role within the frame of NK cell activation.

A recent study has also called into question whether PD-1 is actually expressed at all on NK cells ^37^. Our results are in agreement with some of these findings, including the low expression of PD-1 on NK cells under normal conditions. However, in contrast to the study of Judge *et al* which used IL-2-stimulated NK cells to investigate PD-1 expression on NK cells^37^, we established here that the combination of IL-12, IL-15 and IL-18 leads to PD-1 expression, which is in line with previous publications ^13,57,58^. Furthermore, we did not see much expression of PD-1 on splenocytes but we could see that there was clear expression of PD-1 on tumor infiltrating particularly since we used *PD-1*-deficient mice as controls. When comparing our two tumor models, PD-1 expression on NK cells seemed to be highest when infiltrating the MTAP1A tumors rather than RMA-S. The MTAP1A tumor was generated by painting the skin with methylcholanthrene whereas RMA-S is lymphoma. Therefore, the tumor microenvironment (TME) within these tumors might determine the level of expression of PD-1. For example, fibrotic tumors have been associated with TGF-*β* ^59^ and TGF-*β* itself can induce PD-1 expression on T cells^60^. This suggests that the microenvironment surrounding NK cells could lead to PD-1 expression. Metzger *et al.* have also suggested that false positives can be obtained by anti-PD-1 antibodies binding to nuclear antigen in dying cells ^61^. In our studies, we have compared our staining of WT NK cells with NK cells from *PD-1*^−^/^−^ mice and did not see non-specific binding using the anti-PD-1 antibody clone <u>RMP1-14 nor clone 29F.1A12</u>. We did not use the J43 clone in our studies as we had previously found some non-specific binding with this clone. This suggested that at least in our hands, our observed PD-1 expression was not due to cross-reactivity with another antigen. Finally, since PD-1 is expressed on other tumor-infiltrating cells, there is still the possibility that PD-1 may be transferred by trogocytosis from surrounding cells to NK cells via PD-L1. We cannot rule this out but since we see PD-1 expression by mRNA from NK cells, this probably suggests that the observed increase in PD-1 expressing NK cells is at least in part through induction of PD-1.

NK cells play an important role in clearance of tumor cells, and impairment of NK cell functions results in an increased risk for the development of cancer. Both tumor-infiltrating NK cells (TINKs) and tumor-associated NK cells (TANKs) have been described^62^, but their function and expression profiles have yet to be defined. Our single cell gene expression data reveal that NK cells within the tumor microenvironment (<u>TME</u>) separate into five distinct clusters. Many DE genes of our intra-tumoral NK cells have been previously described in tissue resident NK cells in different organs including liver, lung, lymph node and placenta. The high expression of tissue residency markers in NK cells within the TME could indicate that these NK cells are tumor tissue resident. Whether these NK cells infiltrate tumors (TINKs) to eliminate them, or whether they associate with tumor cells (TANKs) and facilitate pro-angiogenic properties, remains difficult to assess. In many tumors, TINKs exhibit a profoundly altered phenotype with defects in degranulation and IFNγ expression^63^. It is still unclear whether PD-1 on NK cells leads to exhaustion and functional impairment, or if expression of PD-1 restricts NK cell activation to prevent the exhausted phenotype, as has been suggested for T cells^24,48^.

Previous work has established the bidirectional signaling of PD-1 and PD-L1. *Cis* interactions between PD-1 and PD-L1 on antigen presenting cells have been shown to decrease availability of PD-L1 for *trans* binding to PD-1 on T cells, and both *cis* and *trans* interactions are susceptible to antibody blockade ^46^. In the current study, we have shown that *cis* interaction between PD-1 and PD-L1 and a potential sequestration of available PD-1 for *trans* signaling also occurs on NK cells. We show that in the absence of PD-1, the diffusion rate of PD-L1 is significantly increased, while the size of PD-L1 clusters is decreased, indicating that PD-L1 forms clusters with PD-1 on the same membrane, thus limiting the movement of PD-L1 and potentially also that of PD-1. This suggests that the levels of PD-L1 on NK cells can determine their response to PD-1 signaling imposed by PD-L1^+^ cells inside the tumor microenvironment. We further provide a model for how PD-1 and PD-L1 interact in *trans* and in *cis*, where the same amino acid residues are involved in these interactions.

The binary PD-1/PD-L1 complex was crystallized both for human PD-L1 and murine PD-1 ^64^ and for human PD-L1 and human PD-1 ^65^. In both cases, protein-protein binding occurs via “cheek to cheek interaction” of Ig domains of PD-1 and PD-L1, and this was almost identical in the two structures. We hypothesize that the long flexible stalk of PD-1 allows both *cis* and *trans* interaction, where PD-1 “tip-toes” to reach PD-L1 with an extended stalk, while keeping the same PD-1/PD-L1 “cheek-to-cheek” interface found in the crystal structures. The stalk region of PD-1 (residues R147-V170) was modelled in extended conformation to demonstrate that its length is sufficient to allow both *cis*- and *trans-*interaction with the N-terminal domain of PD-L1. High sequence homology between murine and human proteins (77% for the PD-L1 and 64% for PD-1) and conservation of the residues forming intermolecular hydrogen bonds suggest that the *cis* and *trans*-interaction for the PD-1 and PD-L1 could be possible for the human cells as well. Indeed, *cis* binding of human PD-1 and human PD-L1 has recently been demonstrated ^46^.

A recent study has shown that NK cells up-regulate PD-L1 in response to IFN-γ and that NK cells from AML patients show increased expression of PD-L1 ^66^. PD-L1^+^ cells in the TME negatively regulate PD-1^+^ effector cells, but at the same time, PD-L1 on T and NK cells might inhibit survival of PD-1^+^ APCs ^67^. In addition to binding PD-1 in *cis*, PD-L1 can also bind to CD80 on the same membrane, which may repress both PD-1 and CTLA-4 signaling while favouring the CD28 axis ^68^. These multi-facetted binding patterns in *trans* and *cis* may contribute to the fine tuning of the immune response within the TME, and may be the cause for the differences observed when treating cancer patients with anti-PD-1 vs. anti-PD-L1 blocking antibodies ^69^.

Antibody immunotherapy against PD-1 or its ligands has emerged as one of the breakthrough immunotherapies in the clinics ^70^. PD-1 was expressed on NK cells in a number of clinical studies examining patients with different cancers ^11,14,15,49^. NK cells expressing PD-1 in human cancers appear to be confined to more mature NK cells ^15^. However no data exists on the effects of anti-PD-1/PD-L1 therapy has on NK cell function and development. Within the mouse NK cells, we did not find to date a specific NK cell population/subset that expressed PD-1. However it may well be that anti-PD-1 therapy does not only affect adaptive cells and also innate cells that would have potential knock-on effects on adaptive immune-responses ^71^.

## MATERIAL AND METHODS

### Mice

C57BL/6, *PDCD-1*(PD-1)^−/−^ (generously provided Dr. Tasuku Honjo, Kyoto University, Kyoto, Japan) ^26^, *RAG1*^−/−72^ and *PD-1*^−/−^x*RAG1*^−/−^ (PD-1xRAG1^−/−^) ^73^, *H-2K*^*b*^*xH-2D*^*b*−/−^ (MHC-I^−/−^) ^74^ mice on the C57BL/6 background were housed under specific pathogen free conditions at the Department of Microbiology, Tumor and Cell Biology and Astrid Fagraeus Laboratories, Karolinska Institutet, Stockholm. All procedures were performed under both institutional and national guidelines (Ethical numbers from Stockholm County Council N147/15). Sex and aged match mice were used for all experiments. Mice were chosen randomly for control or treated groups.

### Tumors

MHC-I-deficient lymphomas RMA-S (*TAP2*-deficient), and *TAP1*-deficient MCA fibrosarcoma (clone MTAP1A) have been previously described ^6,36^. RMA-S cells were inoculated at the LD_50_ dose of 10^5^ s.c. in the flank of mice. MTAP1A was inoculated at a dose of 10^5^ cells/mouse. Tumor growth was measured every two days and mice were sacrificed when the tumor reached 10^3^ mm.

### NK cell purification and culture

Single-cell suspension from spleens was depleted of erythrocytes, and NK cells were positively sorted using anti-DX5^+^ magnetic beads or by negative sorting using MACS separation, according to the manufacturer’s instructions (Miltenyi Biotec, Bergisch Gladbach, Germany). Cells were resuspended in complete medium (αMEM; 10 mM HEPES, 2 × 10^−5^ M 2-ME, 10% FCS, 100 U/ml penicillin, 100 U/ml streptomycin) with with 100 ng/ml mouse IL-12 (PeproTech), IL-15 (Immunotools) and 100 ng mouse IL-18 (MBL International, Woburn, MA, USA) for 4 days. For isolation of NK cell subsets, NK cells were isolated as above and then sorted on MoFlo XDP cell sorter (Beckman Coulter, Brea, CA, USA).

### In vivo rejection assay

Splenocytes from B6 or MHC-I^−/−^ mice were labeled with 0.5 μM CFSE (target cells) or 0.5 μM CellTrace Violet (control cells; Thermo Fisher Scientific Life Sciences) for 10 min. Target and control cells were washed, then mixed and 1–3 × 10^6^ cells coinjected intravenously via the tail vein into B6, PD-1^−/−^ mice or MHC-I^−/−^ mice as controls for NK cell-mediated killing. The injection mix was analyzed by flow cytometry for reference. Two days later, the spleens were harvested and erythrocytes depleted, and the relative percentages of target and control cells were measured by flow cytometry ^31^. Rejection was estimated as the relative survival of target or cells, calculated as: % remaining target cells of labeled cells/% target cells in inoculate or % remaining control cells of labeled cells/% control cells in inoculate.

### Antibodies and Flow Cytometry

Antibodies used in the study were purchased from BD, Biolegend or eBioscience. Clones for the different antibodies were: anti-NK1.1 (clone PK136 Biolegend), -NKp46 (29A1.4), -PD-1 (RMP1-14 and 29F.1A12), -NKG2A (20d5 Biolegend), Ly49A (YEI/48 BD) Biolegend), Ly49C (5E6), Ly49D (4E5), Ly49G2 (4D11), Ly49H (3D10 eBioscience), Ly49I (YLI-90, eBioscience) -CD11b (clone M1/70, BioLegend), -CD127 (clone A7R34, BioLegend), -GITR (DTA-1, Biolegend), -CD244 (m2B4(B6)458.1), -TIGIT (GIGD7, eBioscience), CD39 (Duha59, Biolegend), -KLRG1 (clone MAFA, Biolegend), -CD226 (10E5, Biolegend),-GR1 (GR1, Biolegend),- CXCR3 (CXCR3-173 eBioscience), -CXCR4 (clone L276F12 Biolegend)-CD274 (clone MIH1, Biolegend). The 4LO3311 (Ly49C) hybridoma was a kind gift from Suzanne Lemieux.

Flow cytometry was performed on CyAN ADP LX 9-colour flow cytometer (Beckman Coulter, Pasadena, CA) or LSRII (Becton Dickinson). Data were analyzed using FlowJo software (Tree Star Inc, OR).

### Molecular modelling of cis- and trans-interactions between PD-1 and PD-L1

Three-dimensional molecular models of the full-length extracellular regions of murine or human PD-1/PD-L1 complexes (PD-L1 residues 19-239 and PD-1 residues 21-170) were created based on the crystal structure of the chimeric complex of human PD-L1 and murine PD-1 (pdb code3BIK)^64^. To our knowledge, no crystal structure of murine PD-L1 has been determined yet, although several crystal structures of human PD-L1 are available ^64,75^. The crystal structure of human PD-L1 revealed that it consists of two Ig domains linked by a 10 residues-long stalk region. The sequence identity between murine and human PD-L1 is 77%, which means that their 3D structures may be very similar. Indeed, the model of murine PD-L1 created using SwissModel^76^ is very similar to human PD-L1. Replacement of human PD-L1 with its murine orthologue in the 3BIK structure allowed us to generate a full-length model of the murine PD-1/PD-L1 complex. Conversely, replacement of murine PD-1 with the human orthologue allowed us to create a three-dimensional model of the full-length human PD-1/PD-L1 complex. The stalk regions of PD-1 (residues 147-170) and PD-L1 (residues 229-239) were modelled in an arbitrary extended conformation using the program Coot^77^ followed by model regularization to improve the geometry of the peptide chain and remove all possible sterical clashes.

## SMART-SEQ2

scRNA-Seq was performed in 384-well format. The tumors were isolated, rapidly processed, stained for a panel of surface markers and single cell sorted within approximately 90 minutes of organ harvest. In total 382 NK cells were sorted directly into 2 μl lysis buffer using a BD Influx from pooled tumors from either 3 WT and 3 PD-1^−/−^ KO mouse respectively. SMART-Seq2 libraries were prepared using the method described in Picelli et al. ^38^ by the Eukaryotic Single Cell Genomics national facility at SciLife Laboratory, Stockholm.

Digital gene expression matrices were preprocessed and filtered using the Seurat v3.0 R package (https://github.com/satijalab/seurat). Outlier cells were first identified based on 3 metrics (library size, number of expressed genes, etc). Low abundance genes were removed by removing all genes that were expressed in less than 3 cells. The raw counts were normalized and transformed using the ‘LogNormalize’ function of Seurat. Highly variable genes were detected using the proposed workflow of the Seurat R package. Unsupervised clustering of the cells was performed and visualized in two-dimensional scatterplots via Uniform Manifold Projection (UMAP) function using the Seurat R package.

### Microscopy and FCS analysis

#### Diffusion of PD1 and PDL1 on cell surface

Zeiss 510 microscope with a Confocor 3 system (Carl Zeiss Microimaging GmbH), C-Apochromat 40x/1.2 NA water objective was used for Fluorescence Correlation Spectroscopy (FCS)measurements ^78^. Diffusion of interested molecules were measured using fluorescent labelled antibodies and FCS measurements were calibrated by measuring Alexa-488 and Alexa-647 dyes in solution at different power scale concentration whose diffusion coefficient is known. For cell preparation, spleens were isolated from from *RAG1*^−/−^ and *PD-1xRAG1*^−/−^ mice. From single cell suspension of splenocytes of mice, NK cells were isolated by MACS NK cell isolation kit mouse II (Miltenyi Biotech Norden AB, Sweden). NK cells were stained for PD-1-Alexa flour 488 and PD-L1-Alexa flour 647, and microscopic chambers were coated with poly-L-lysine, so the cells are made to attach to the glass surface ^45,79^. All the FCS measurements on cells were made on the cell surface for the diffusion of PD-1 and PD-L1.

#### FCS analysis

FCS Data was analyzed using MATLAB based written algorithm to have graphical user interface (GUI) for fitting. GUI permits to assume the initial fit coefficient like N-number of molecules, Tau D-Diffusion time for the molecule to diffuse within the focal volume, triplet state of the molecules. Different fit models and time fit domain was considered for free dyes and cells. Where 3D diffusion model fit was chosen for free dyes with time domain fit 0.5 μsecond to 0.1 msecond and 2D diffusion model fit for cells with time fit between 1 millisecond to 5 second.

### Statistical analyses

All statistical analysis was performed using GraphPad Prism software (La Jolla, CA).

## Data availability

Smart-Seq2 data will be made available at upon acceptance. Other data is available from the corresponding author upon reasonable request.

## Author Contributions

Conceptualization: A.K.W., N.K., C.T., B.J.C.; methodology: A.K.W., N.K., C.T., S.B.S., P.R., T.S., A.A., K.K., B.J.C; Data collection: A.K.W., N.K., C.T., K.v.d.V., S.B.S., D.O., E.LG., N.C., S.T., T.S., B.J.C;; Analysis and interpretation: A.K.W., N.K., C.T., K.v.d.V., S.B.S., D.O., E.LG., N.C., S.T., P.R., T.S., A.A., K.K., B.J.C.; writing - original draft preparation, A.K.W., C.T. T.S., B.J.C.; critical revision of the article: A.K.W., N.K., A.A., K.K., B.J.C., visualization, A.K.W.,C.T., S.B.S., T.S., B.J.C; funding acquisition, K.K. and B.J.C.; All authors have read and agreed to the published version of the manuscript.

## Conflicts of Interest

The authors declare no conflict of interest.

## Acknowledgement

ScRNA-seq was performed at the eukaryotic single-cell genomics facility at SciLife laboratories (Stockholm, Sweden). The data handling was enabled by resources provided by the Swedish National Infrastructure for Computing (SNIC) at Uppsala partially funded by the Swedish Research Council through grant agreement no. 2018-05973. We also thank Jonas Søndergaard for advice regarding the scRNA-seq analyses.

## Notes

Financial Support, This work was funded by the Swedish Cancer Society, Swedish Research Council, the Karolinska Institute Foundations and the Swedish Foundation for Strategic Research.

None of the authors have any conflicts of interest.

### Competing Interest Statement

The authors have declared no competing interest.

