## Supplemental Figures and Table for "PD-1 expression on NK cells can be related to cytokine stimulation and tissue residency"

### Supplemental Information

#### Subsets of NK cells express PD-1 upon cytokine stimulation and tissue residency

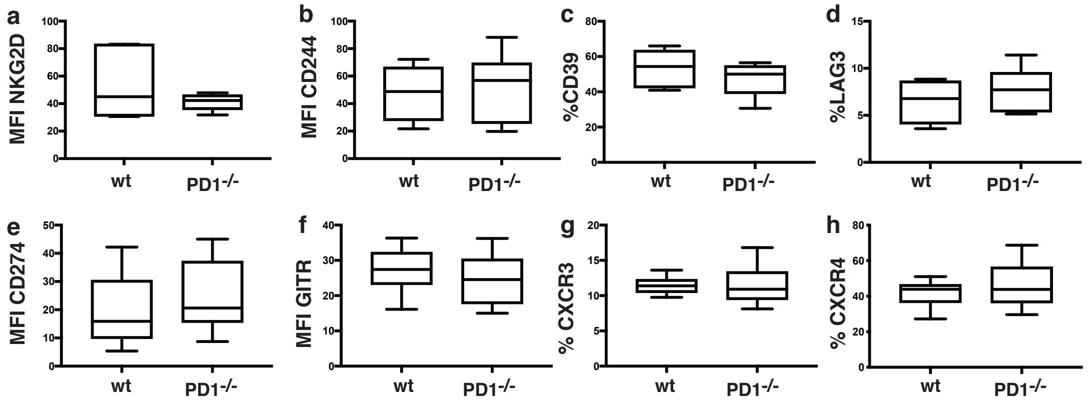


**Supplemental Figure 1. Expression of NK cell markers and chemokine receptors on wt and PD-1$^{-/-}$ NK cells.** (a) Percent of NK cells expressing CD39, (b) LAG3, (c) MFI of CD244, (d) MFI of GITR, (e) Percent of NK cells expressing CXCR3 (f) Percent of NK cells expressing CXCR4 (g) MFI of CD274.

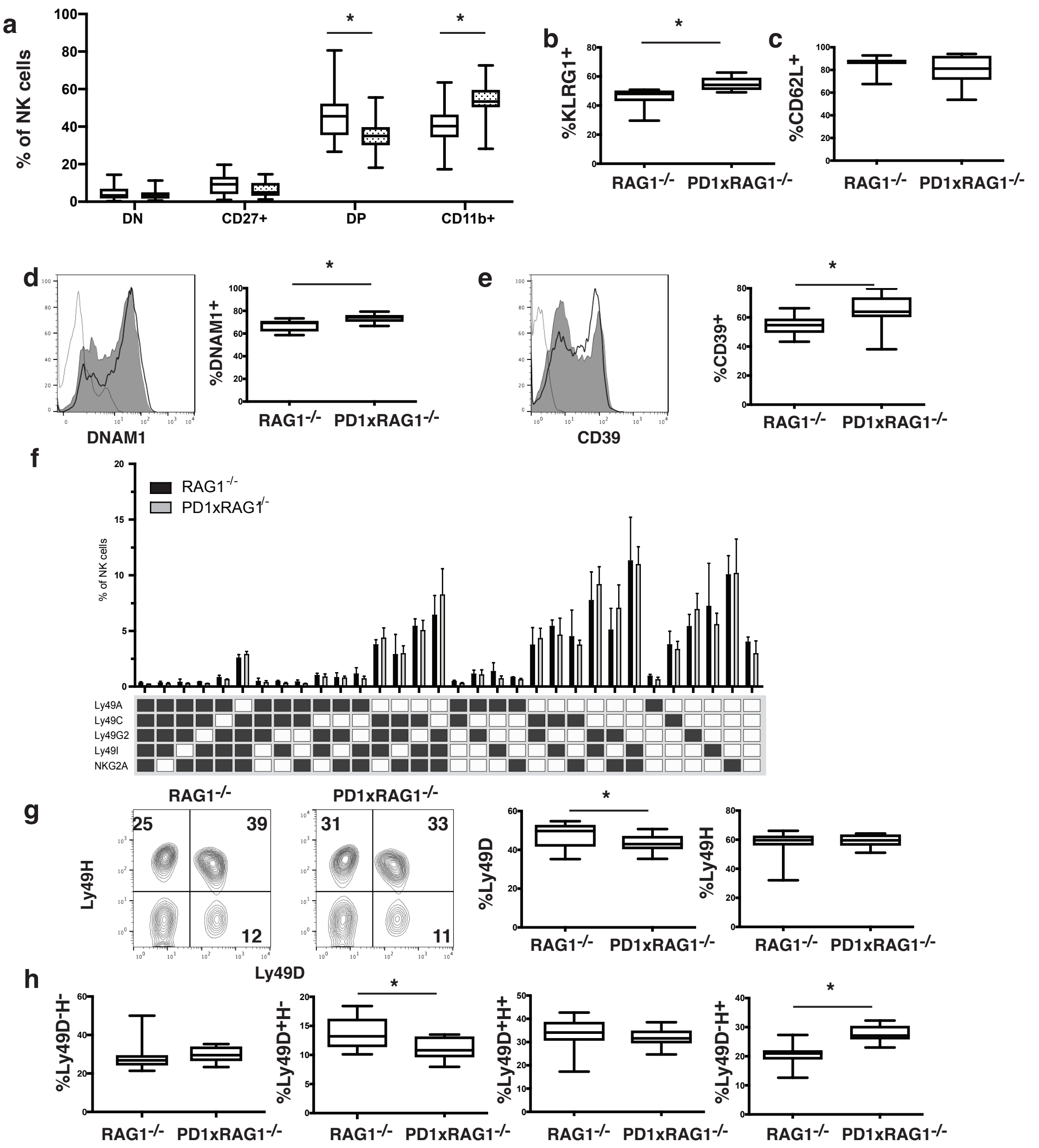


**Supplemental Figure 2. Phenotype of NK cells from RAG1$^{-/-}$ and RAG1$^{-/-}$PD-1$^{-/-}$ mice.** (a) Expression of CD11b and CD27 on NK cells from WT and PD-1$^{-/-}$ mice (*p<0.01 MannWhitney test, n=18-20 mice. (b) Expression of KLRG1 on NK cells from WT and PD-1$^{-/-}$ mice (*p<0.01 MannWhitney test, n=18-20 mice). (c) Expression of CD62L on NK cells from WT and PD-1$^{-/-}$ mice. (d) Expression of DNAM-1 on NK cells from WT and PD-1$^{-/-}$ mice, bar graphs represent percent expressing cells and the mean floursecent intensity of expression (*p<0.01 MannWhitney test, n=18-20 mice. (e) Expression of inhibitory Ly49 molecules and NKG2A on NK cells from WT and PD-1$^{-/-}$ mice (*p<0.01 Mann Whitney). (f) Expression of activating Ly49 molecules on NK cells from WT and PD-1$^{-/-}$ mice. (*p<0.01 MannWhitney test, n=18-20 mice (g) Expression of Ly49D and Ly49H populations on NK cells from WT and PD-1$^{-/-}$ mice (*p<0.01 MannWhitney test, n=18-20 mice).

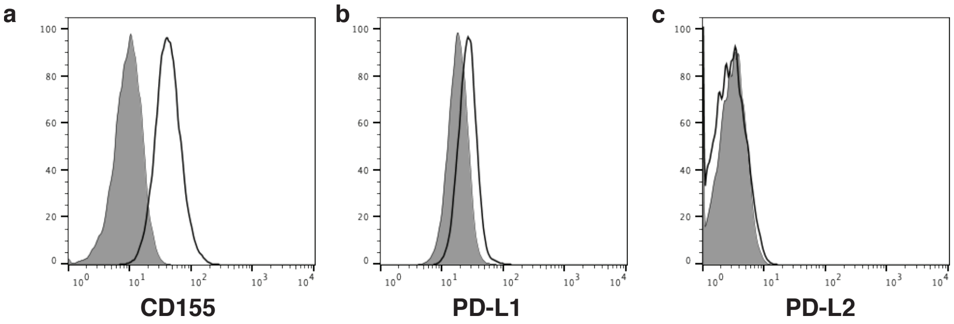


**Supplemental Figure 3. Expression of CD155, PD-L1 and PD-L2 on MTAP1A tumor cells.** MTAP1A tumor cells were stained with mAbs specific for CD155 (a), PD-L1 (b), or PD-L2 (c) and analyzed by Flow cytometry for expression of these markers.

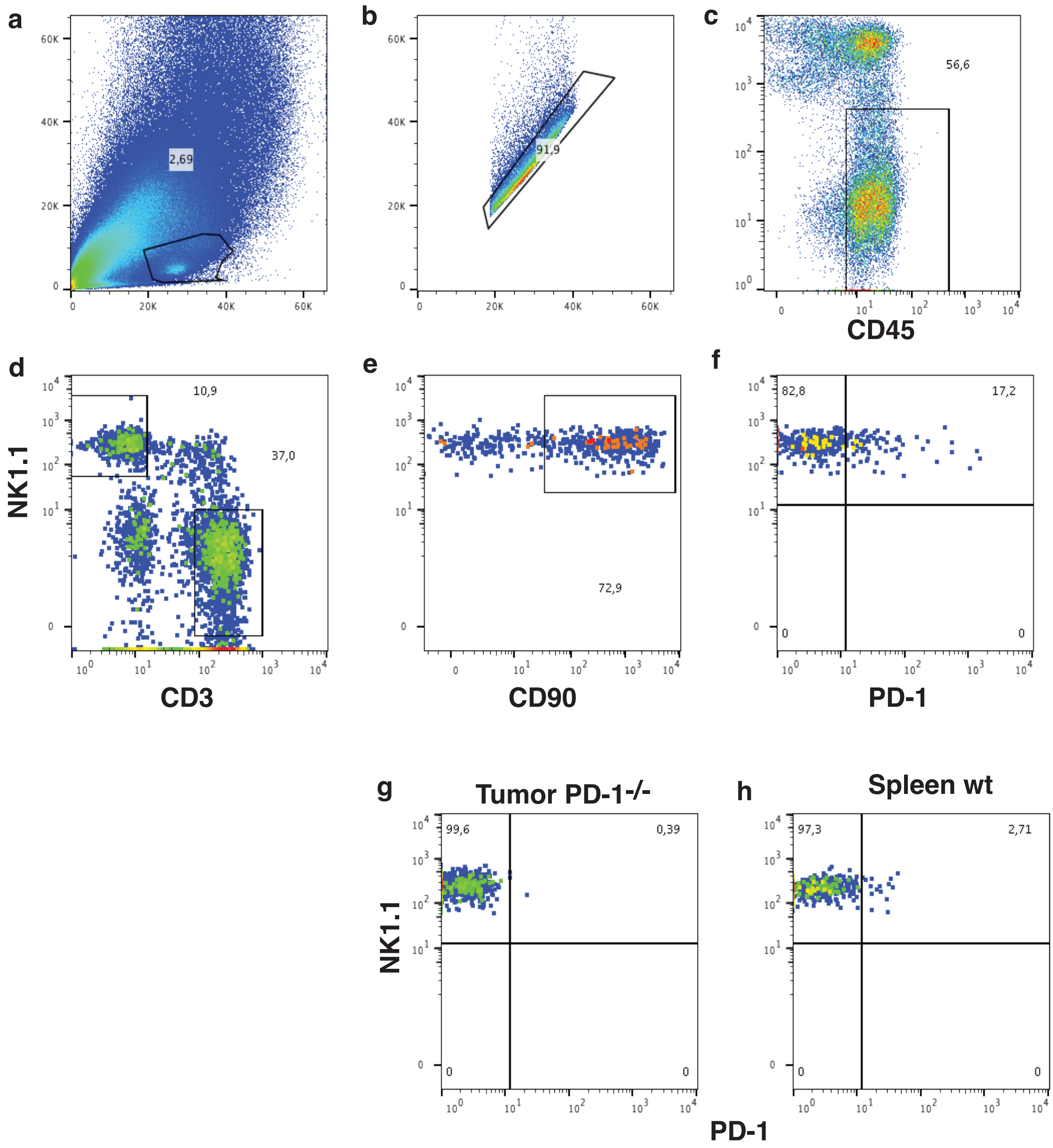


**Supplemental Figure 4. Gating strategy for intratumoral NK cells.** MTAP1A tumors were excised, crushed and filtered to get a single cell suspension. Flow cytometric analyses (a-b) of markers for leukocytes (c), NK cells and CD3 (d-f) were performed. PD-1 expression was assessed on intratumoral NK cells in PD1-ko mice (g) and spleen NK cells of wt mice (h).

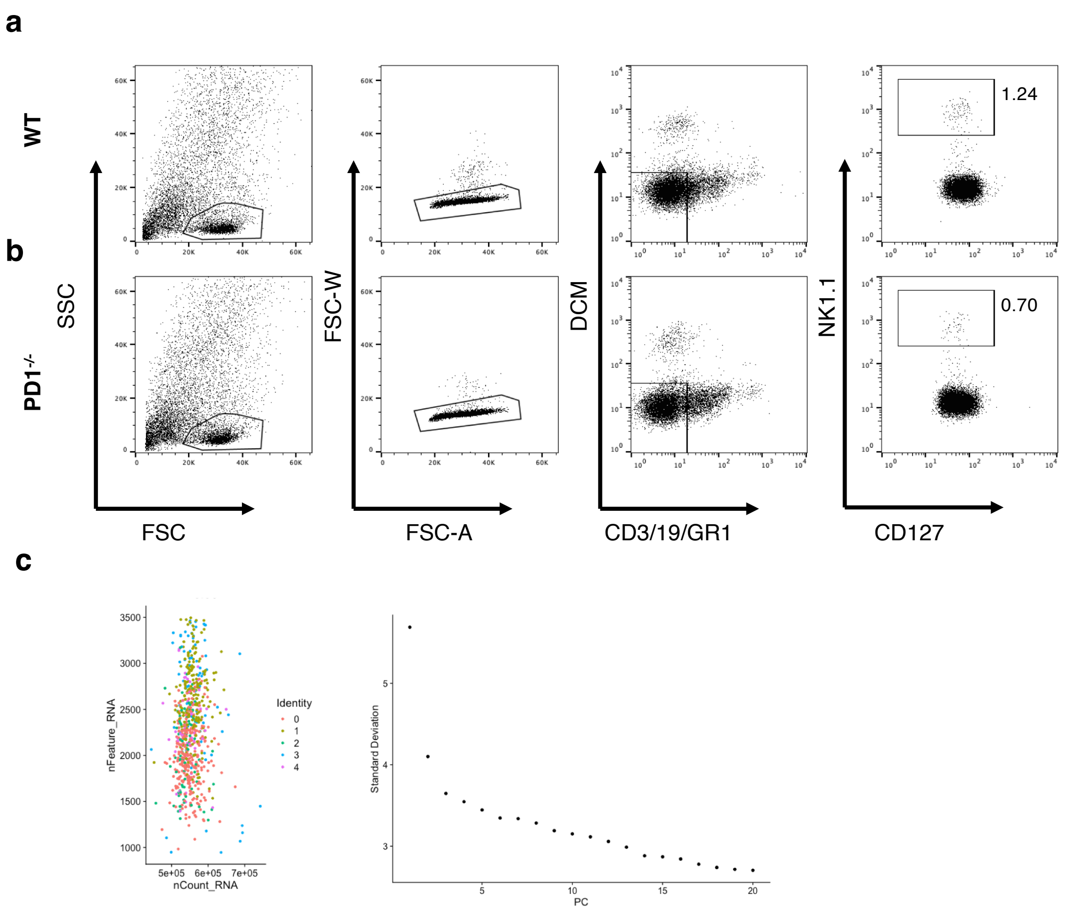


**Supplemental Figure 5. Sorting strategy and scRNA-SEQ QC.** Viable NK cells (NK1.1$^+$CD3$^-$CD19$^-$GR1$^-$) were sorted from pooled tumors from either WT (a) or *PD1*$^{-/-}$ (b) according to the gating strategy shown. (c) SMART-SEQ2 data was processed according to the standard Seurat v3 pipeline with an elbow plot used to determine cutoff for dimensionality reduction.

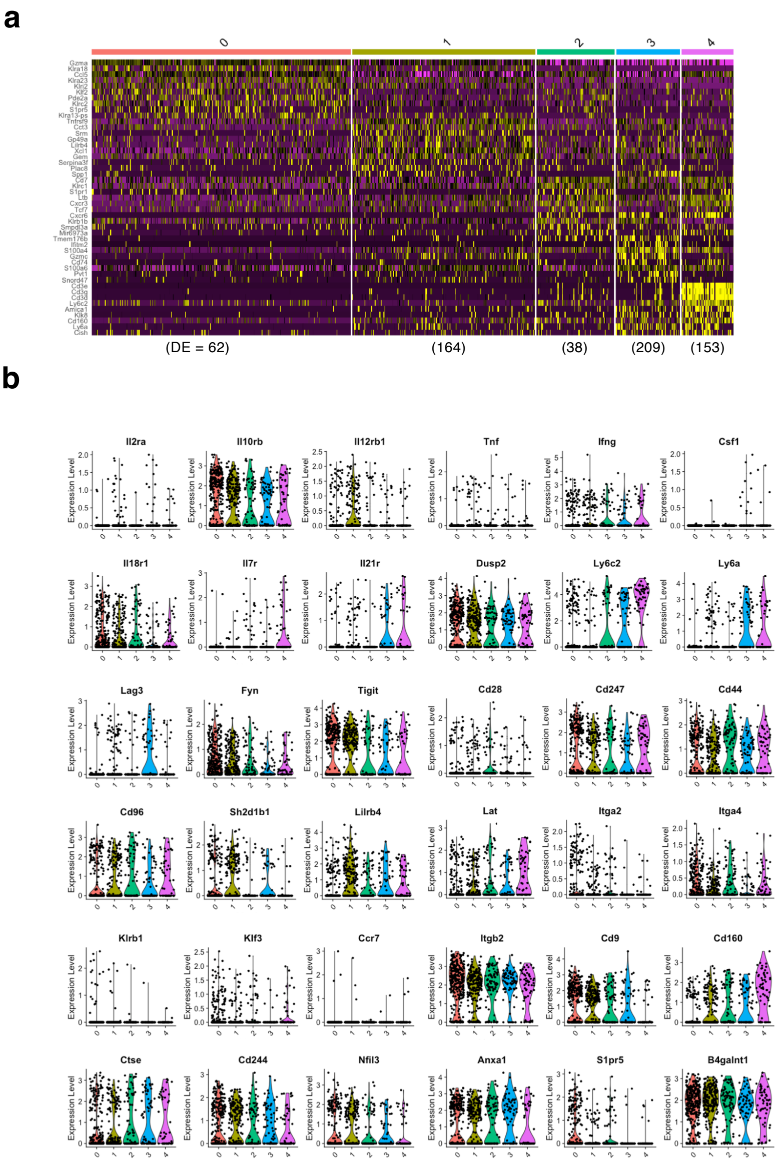


**Supplemental Figure 6. Extended analysis of scRNA-Seq of infiltrating WT and *PD1*$^{-/-}$ NK.** (a) Heatmap visulation of top 10 genes per. Number of DE genes per cluster is shown below. (b) Violin plots showing expression of the genes Il2ra, Il10rb, Il12rb1, Tnf, Ifng, Csf1, Il18r1, Il7r, Il21r, CD44, Ly6c2, Ly6a, Lag3, Fyn, Tigit, CD28, CD247, CD96, Sh2d1b1, Lilrb4, Lat, Itga2, Itga4, Klrb1, Klf3, Ccr7, Itgb2, Cd9, CD160, CD5, CD244, Nfil3, Anxa1, S1pr5 and B4galnt1, where expression significantly differs between WT and PD-1$^{-/-}$ NK cells.

**Supplemental Table 1: P values of genes that differ in their expression between WT and PD-1-KO NK cells.**

| Gene name | p_value |
|---|---|
| Serinc3 | 6.27677503191847e-31 |
| Hist1h4m | 6.55865752557258e-23 |
| Ctse | 1.15768218666942e-15 |
| Hist1h1e | 1.11246146661204e-09 |
| Wdfy1 | 1.06702627915021e-08 |
| Ugt1a7c | 1.95929659888963e-08 |
| Cxcr6 | 1.97343268509504e-07 |
| Cd226 | 3.13682908855125e-07 |
| Ramp1 | 9.29433013791078e-07 |
| Lgals1 | 1.80274751617846e-06 |
| Nedd9 | 5.00263546868797e-06 |
| Fgl2 | 1.13557269866248e-05 |
| Plac8 | 2.6879413217007e-05 |
| Ms4a4c | 3.89740981356844e-05 |
| Mapk14 | 4.34626898068145e-05 |
| Ly6c2 | 9.21126370197399e-05 |
| Ctsz | 0.000110090159134527 |
| Aarsd1 | 0.00014546275752589 |
| Cndp2 | 0.000168918497256701 |
| Ly6a | 0.000181160676017526 |
| Klrb1b | 0.000215424070178674 |
| Dusp2 | 0.000255515137305345 |
| Gyg | 0.000336207620171877 |
| 4930486L24Rik | 0.000400511875558075 |
| Vmp1 | 0.00041638049673073 |
| Mcm6 | 0.000466756919579111 |
| Capg | 0.000754405873530379 |
| Cd300lf | 0.000806216994065812 |
| Klrc1 | 0.000906898344562637 |
| Lgals3 | 0.000955605594016018 |
| Ccl4 | 0.00169530836212815 |
| Stk25 | 0.00219701812850478 |
| Zfp36 | 0.00221040436852507 |
| Styk1 | 0.00256774322556803 |
| Cd160 | 0.00376332357863304 |
| Trappc2l | 0.00383832466555109 |
| Nfkb2 | 0.00384483049748892 |
| Tmem199 | 0.00410912503961128 |

| | |
|---|---|
| Lag3 | 0.00414929516522654 |
| Ifi27 | 0.00474219709535523 |
| Cysltr2 | 0.0051475976240368 |
| Eif2d | 0.00525748204514534 |
| Klrg1 | 0.00898488940611906 |
| Glrx5 | 0.0228145183424616 |
| Tmem176b | 0.0303113229640398 |
| Klra21 | 0.0307030769116372 |
| Mgst2 | 0.0307433799133878 |
| Mmadhc | 0.0378199706460093 |
| 2900060B14Rik | 0.0404041545808012 |
| Cd3e | 0.0450593340578062 |
| Ifng | 0.062829998418396 |
| Klhl28 | 0.0678278262427947 |
| Cd3g | 0.0719380240787541 |
| Hilpda | 0.0727772544214674 |
